# Pgc-1α is an exercise-responsive regulator of myosteatosis in older individuals

**DOI:** 10.1101/2023.02.07.527478

**Authors:** Hirotaka Iijima, Fabrisia Ambrosio, Yusuke Matsui

**Author notes:** Corresponding and lead contact author: Hirotaka Iijima, PhD, PT, Furo-cho, Chikusa-ku, Nagoya, 464-8601, Japan. Co-corresponding author: Yusuke Matsui, PhD, 1-1-20 Daiko-Minami, Higashi-ku, Nagoya, 461-0047, Japan.

## Abstract

Accumulated fat in skeletal muscle, common in sedentary older individuals, compromises skeletal muscle health and function. Mechanistic understanding of *how* physical activity levels dictate fat accumulation represents a critical step towards establishment of therapies that promote healthy aging. Using a network paradigm that characterized the transcriptomic response of aged muscle to exercise versus immobilization protocols, this study uncovered a novel molecular cascade that regulates the fate of fibro-adipogenic progenitors (FAPs), the cell population primarily responsible for the fat accumulation. Specifically, gene set enrichment analyses (GSEA) with network propagation revealed *Pgc1*α as a functional hub of a large gene regulatory network underlying the regulation of FAPs by physical activity. Integrated *in silico* and *in situ* approaches to induce *Pgc-1*α overexpression promoted mitochondrial fatty acid oxidation and inhibited FAPs adipogenesis. These findings suggest that *Pgc1*α is a master regulator by which physical activity regulates fat accumulation in aged skeletal muscle.

**GRAPHICAL ABSTRUCT:** 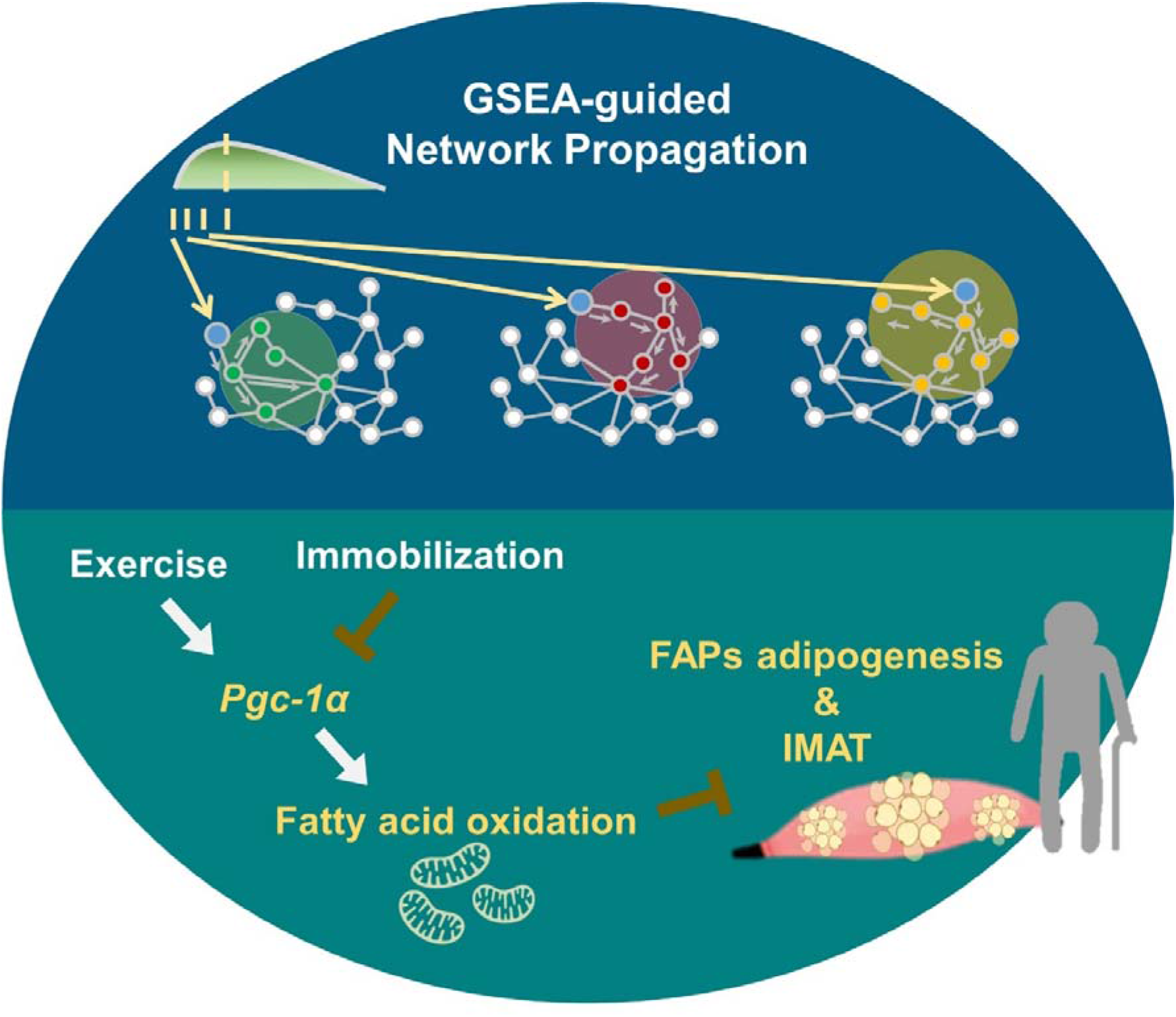

## INTRODUCTION

A unique feature of skeletal muscle aging is accumulation of intra- and intermuscular adipose tissue (IMAT), defined as an ectopic fat found beneath the fascia and within the muscles^1^. Despite the fact that IMAT accounts for only a small portion of total body fat, it is considered a leading cause of insulin resistance and decreased muscle strength^2,3^. IMAT has even been used as a predictor of all-cause and cardiovascular mortality in elderly^4^. As such, there is great interest in better understanding the mechanisms of age-related IMAT to guide the development of therapeutic intervention as well as prognostic biomarkers in a geriatric population.

Accumulation of IMAT is driven, at least in part, by muscle-resident fibro-adipogenic progenitors (FAPs), which are identified by expression of platelet-derived growth factor receptor alpha (PDGFRα)^5-7^. While FAPs are key regulators of muscle regeneration and homeostasis, they also contribute to chronic inflammation, fibrosis, and fat deposition when dysregulated^5-7^. Damage to skeletal muscle triggers a transient phase of FAP proliferation that promotes muscle stem cell differentiation, followed by a return to baseline numbers as a result of apoptosis and subsequent phagocytic clearance^8,9^. However, in the setting of chronic muscle inflammation, as is the case with aged muscle^10^, FAP proliferation is maintained at high levels owing to impaired apoptosis and removal of dead cells via phagocytosis^9^. The result is persistence and adipogenic differentiation of FAPs, thereby inducing the buildup of IMAT^5-7^.

IMAT accumulation is believed to coincide with a reduced muscle contractile activity. As such, physical exercise has been proposed as a means to counteract an age-related increase in IMAT and promote skeletal muscle health. Indeed, a recent meta-analysis showed that physical exercise reduces IMAT accumulation in elderly individuals with chronic diseases^11^. On the other hand, immobilization, defined as restricted muscle contractile activity, increases IMAT^12,13^. We found only one study demonstrating that physical exercise promotes senescence in FAPs of healthy young skeletal muscle in mice^14^. As such, molecular and cellular signaling cascades underlying the exercise- and immobilization-driven IMAT regulation under conditions of low grade chronic inflammation remains poorly understood, thereby precluding an establishment of effective rehabilitative strategies to improve skeletal muscle health in an elderly population.

As a first step to address this critical knowledge gap, this study applied integrated computational approaches to a publicly available database of the skeletal muscle transcriptomic response to physical activity and identified a novel candidate driver of the exercise- and immobilization-induced IMAT regulation in aged skeletal muscle. Specifically, we implemented a series of gene set enrichment analyses (GSEA) to the aged skeletal muscle transcriptome and demonstrated that exercise and immobilization differentially regulate FAPs adipogenesis. Next, using a weighted gene co-expression network analysis (WGCNA)^15,16^ integrated with a GSEA-guided squeeze network propagation approach, this study explored possible mechanistic drivers of FAPs adipogenesis. The culmination of these *in silico* analyses suggests that exercise and immobilization regulate *Pgc-1*α and its downstream target, mitochondrial fatty acid oxidation, to regulate FAPs adipogenesis.

## RESULTS

### Resistance exercise and immobilization differentially regulated FAPs adipogenesis genes

While accumulated evidence has shown that IMAT is affected by muscle contractile activity (i.e., IMAT is reduced by resistance exercise but increased by immobilization)^11-13^, the underlying molecular mechanism is unclear. To address this gap, we first summarized the transcriptomic response of elderly skeletal muscle to resistance exercise versus immobilization. For this purpose, we accessed MetaMEx, a publicly database of transcriptomic responses to an array of exercise and immobilization protocols in human skeletal muscle^17^. From 50 studies with 83 data sets curated through the MetaMEx database, our current study finally included 4 studies with 7 data sets for chronic resistance exercise (3 studies with 5 data sets; mean age: 75.9 years old; body mass index: 25.1 kg/m^2^; n = 36 participants)^18-20^ and sedentary (1 study with 2 data sets; mean age: 68.0 years old; body mass index: 25.1 kg/m^2^; n = 18 participants)^21^ (**Figure 1A-B**). In the included studies, healthy elderly participants either completed a resistance exercise protocol 3 days/week for 3-6 months or were immobilized through bed rest for a total of 5 days. All participants in the resistance exercise group had sedentary behaviors at baseline and reported not participating in any formal exercise programs or regular physical activity when enrolling the study^18-20^. Similarly, all participants in sedentary group had minimal levels of physical activity at baseline^21^. In these studies, skeletal muscle tissue was collected from vastus lateralis muscle and used for RNA-seq (Illumina HiSeq) or microarray (Affymetrix) analyses. The integrated transcriptomic data from four studies with seven data sets included 13,589 genes. To account for the bias caused by low expression count data, we applied a filter based on the gene expression proposed by Chen et al^22^ to the original RNA-seq data (GSE97084^23^ and GSE113165^21^). Similarly, to account for the bias caused by low signal intensity data, we applied filtering based on signal intensity^24^ to the original microarray data (GSE28422^25^ and GSE117525^26^). These two different filters finally resulted in 7,468 genes that were used for further analyses.

**Figure 1.**
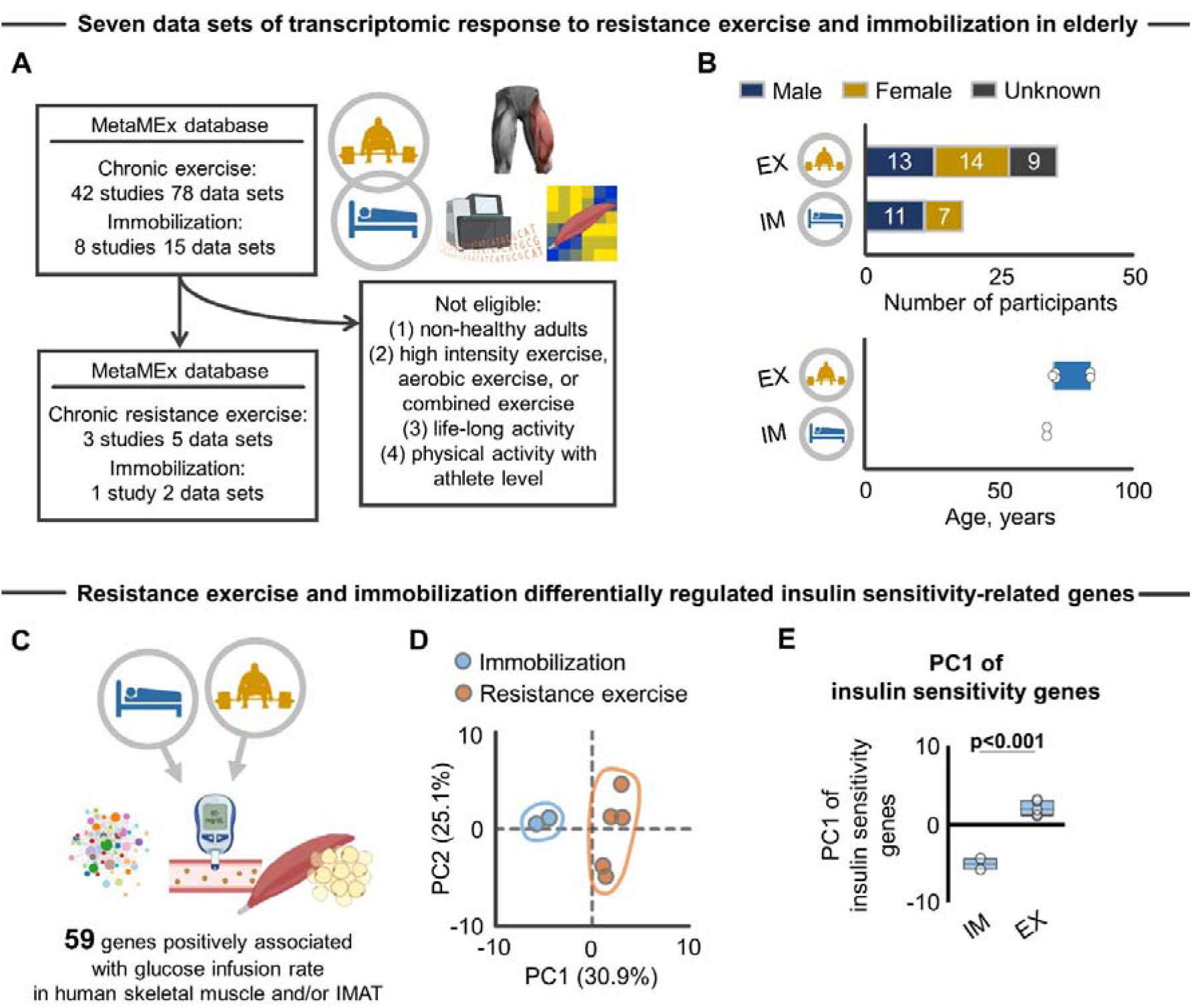
Transcriptomic response of insulin sensitivity-related genes to resistance exercise and immobilization protocols. **A**, Four studies with seven data sets^21,25,26,58^ from MetaMEx database^17^ were included for *in silico* analyses investigating the transcriptomic response of aged skeletal muscle to chronic (≥3 months) resistance exercise (three studies with five data sets) or immobilization (one study with two data sets). **B**, Participants’ characteristics. Each dot represents a single dataset. **C**, Schematic showing the concept of the analysis to characterize transcriptomic response of 59 genes positively associated with glucose infusion rate (i.e., insulin sensitivity) in elderly skeletal muscle. The list of 59 genes were collected from previous literature^27^. **D**, PCA showing the separate clusters in immobilization (blue circle; n = 2) and resistance exercise (orange circle; n = 5) against to overall expression of 59 genes positively associated with glucose infusion rate in elderly skeletal muscle. **E**, PC1 extracted from the PCA revealed a significantly different transcriptomic response of the insulin sensitivity-related genes to immobilization (n = 2) and resistance exercise protocols (n = 5). Statistical analysis was performed using two-tailed Student *t*-test (**E**). Portions of the figures were created with biorender.com. *Abbreviations: EX, exercise; IM, immobilization; PC, principal component; PCA, principal component analysis*.

We first asked whether resistance exercise and/or immobilization changes the expression of genes associated with insulin resistance in skeletal muscle, a primary adverse clinical consequence directly caused by IMAT acccumulation^2^. We accessed a list of 59 genes associated with glucose infusion rate of skeletal muscle (i.e., insulin sensitivity) and/or IMAT in the elderly with obesity and type II diabetes mellitus^27^ (**Figure 1C**). Principal component analysis (PCA) revealed a clear segregation in the transcriptomic response of these insulin sensitivity-related genes when comparing immobilization and resistance exercise protocols (**Figure 1D**). Further, the principal component score, which represents overall gene expression of the insulin sensitivity genes, was significantly different across resistance exercise and immobilization groups (**Figure 1E**). Since the 59 genes were positively associated with glucose infusion rate (i.e., greater insulin sensitivity)^27^, we interpreted this finding to suggest that resistance exercise improved, while immobilization worsened, insulin sensitivity. Opposing effects of resistance exercise and immobilization on insulin sensitivity are well-known clinically^28,29^, thereby indicating the utility of the transcriptomic profile as a reflection of clinical endpoints.

Over the last decade, numerous studies have provided important insights into the cellular origin of IMAT^5-7^. Skeletal muscle contains FAPs that do not form adipocytes under conditions of homeostasis. However, in response to injury, FAPs proliferate and support the commitment of myogenic progenitor cells during muscle repair^30^. The balance between FAPs and myogenic progenitors is, however, disrupted in aged skeletal muscle, owing in large part to chronic inflammation^31^, which drives FAPs to differentiate into adipocytes and give rise to IMAT^5,30,32^. Therefore, we next asked whether exercise and immobilization differentially regulate genes contributing to the adipogenic differentiation of FAPs in aged skeletal muscle. Although no study to our knowledge has identified genes that drive FAPs adipogenic differentiation, Wosczyna et al showed that miR-206 mimicry *in vivo* limits IMAT, while miR-206 inhibition by siRNA recapitulated increased adipogenic differentiation of FAPs *in vitro*^33^. Given that these previous experiments were designed to elucidate the fate switch from FAPs to adipocytes, investigators evaluated gene expression *before* mature adipocytes were formed^33^. We therefore accessed the archived RNA-seq data (GSE171502) that collected RNA from isolated FAPs with and without siRNA to miR-206^33^ (**Figure 2A**). Our RNA-seq analysis identified 161 differentially expressed genes (false discovery rate <0.05) that drive adipocytic differentiation of FAPs (**Figure 2A**). Using these 161 genes, we then performed single sample GSEA (ssGSEA)^34^ to determine whether resistance exercise and immobilization regulate FAPs adipogenesis in opposing directions, as we observed above for insulin sensitivity-related genes (**Figure 1C-D**). GSEA is a computational tool that provides insight into biological processes or pathways underlying a given phenotype^35^. An extension of GSEA^34^, ssGSEA, calculates separate enrichment scores for each paring of a sample and gene set. Results from the two available resistance exercise datasets revealed significant changes in FAP adipogenesis genes, while immobilization significantly changed FAP adipogenesis genes in the opposite direction (**Figure 2B**). We also analyzed male and female data separately to consider sex as a biological variable^36^, and found that male and female participants displayed similar transcriptomic responses to both resistance exercise and immobilization (**Figure 2B**). These results indicate that resistance exercise and immobilization differentially regulate FAPs adipogenesis, independent of sex.

**Figure 2.**
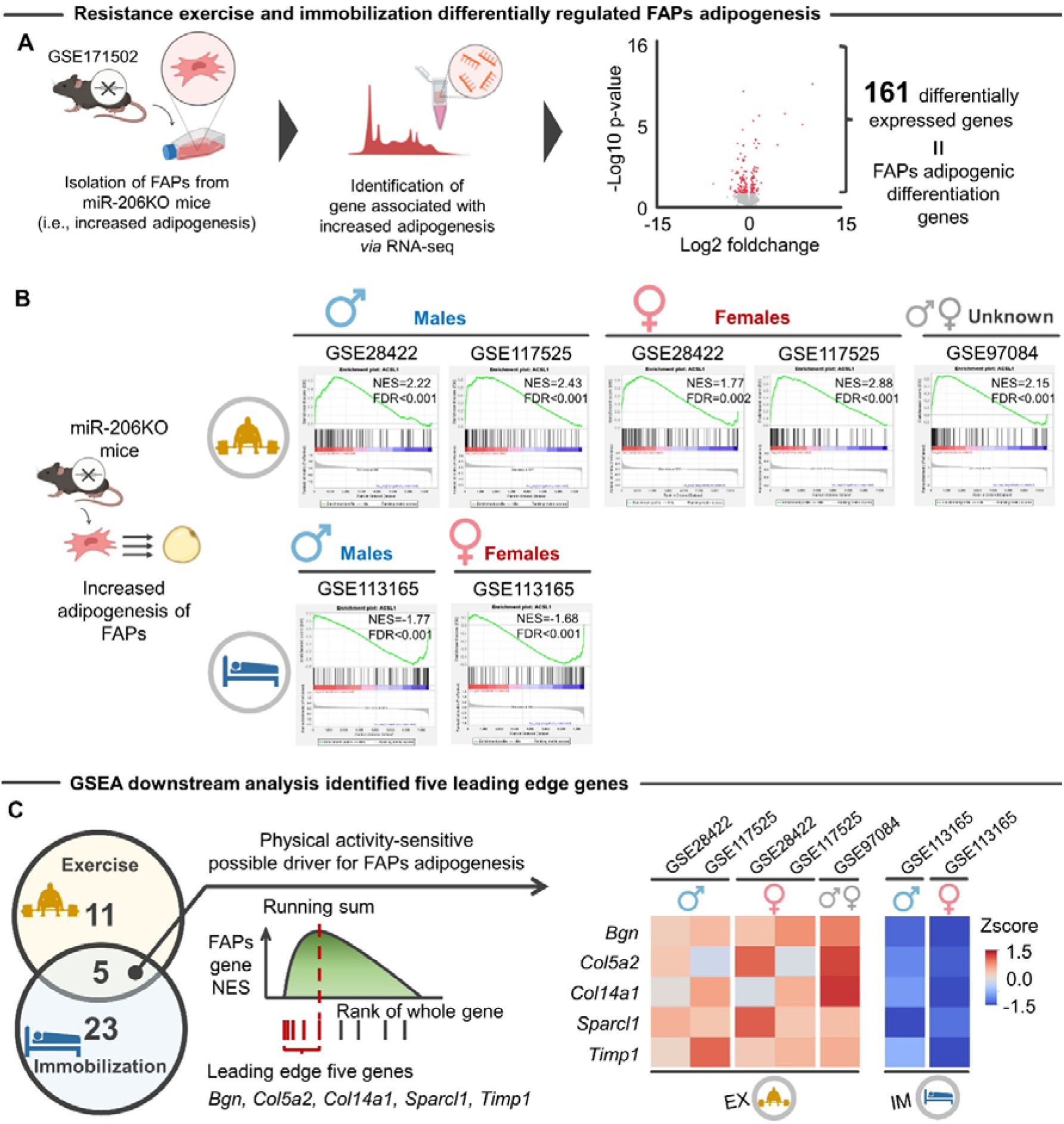
Transcriptomic response of FAPs adipogenesis genes to resistance exercise and immobilization protocols. **A**, Schematic showing the process of defining the genes associated with FAP adipogenic differentiation. From archived RNA-seq of FAPs isolated from the skeletal muscle of wild type and miR-206 KO mice (GSE113165^21^), 161 differentially expressed genes associated with FAPs adipogenesis were identified. **B**, Single sample GSEA revealed that resistance exercise and immobilization regulated FAPs adipogenesis genes in opposing directions, regardless of sex. **C**, Leading edge analysis identified five genes (*Bgn, Col5a2, Col14a1, Sparcl1*, and *Timp1*) that overlapped across exercise and immobilization protocols. Numbers in Venn diagram represent the number of leading edge genes. Color in the heatmap indicates the z-score for each gene. Portions of the figures were created with biorender.com. *Abbreviations: EX, exercise; FAPs, fibro-adipogenic progenitors; FDR, false discovery ratio; IM, immobilization; KO, knock out; NES, normalized enrichment score; GSEA, gene set enrichment analysis*.

To further elaborate the observed relationship between FAP adipogenesis genes and physical activity levels (i.e., exercise vs. immobilization), we next performed leading edge analysis of GSEA, which allows for determination of subsets of genes that contributed the most to the enrichment signal of a given gene set (referred to as the “leading edge subset”)^37^. The leading-edge analysis is determined from the enrichment score, which is defined as the maximum deviation from zero^37^. Genes that comprise a leading-edge subset have a high correlation between their expression level and the phenotype in question (i.e., FAP adipogenesis) and tend to be at the extremes of the distribution, rather than randomly distributed^37^. We found that five leading edge genes (*Bgn, Col5a2, Col14a1, Sparcl1*, and *Timp1*) were significantly changed across the exercise and immobilization protocols, in which resistance exercise upregulated these genes while immobilization downregulated their expression (**Figure 2C**). Among the identified leading-edge genes, *Sparcl1*, a member of the SPARC family, is of particular interest given a recent study demonstrating that *Sparcl1* regulates adipogenesis in mice^38^. *Sparcl1* is a glycoprotein that mediates cell-matrix interactions and is involved in many physiological processes, including cell adhesion, proliferation, differentiation, migration, and maturation^39^. Previous studies have shown that *Sparc* is downregulated with aging but that adeno-associated virus-mediated overexpression of *Sparc* reduced IMAT in a glycerol-injection mouse model^40^. In addition, *Sparc* is a conserved target of miR-29 in skeletal muscle^40^, the latter of which is also regulated by resistance exercise^41^. Taken together, the findings suggest that FAP adipogenesis is suppressed by resistance exercise but exacerbated by immobilization, and that this effect is due, at least in part, to differential regulation of *Sparcl1*.

### WGCNA identified a *Pgc-1*α-centered module associated with FAP adipogenesis genes that are regulated by exercise and immobilization

We next sought to identify mechanistic drivers of FAP adipogenesis that are differentially regulated by resistance exercise and immobilization. Of the five identified genes, only *Sparcl1* has been investigated in the context of skeletal muscle adipogenesis, while the role of the other four leading-edge genes (*Bgn, Col5a2, Col14a1*, and *Timp1*) remains unexplored. As such our interest was directed to other genes that have similar function with the identified leading-edge genes with the aim of obtaining further mechanistic insight into how physical activity levels regulate FAPs adipogenesis. Network medicine-based evidence states that when a gene or molecule is implicated as having a pathogenic role, its direct interactors are likely to play a role in the same pathological process^42^. According to this “disease module hypothesis”, possible mechanistic drivers of exercise- and immobilization-induced FAP adipogenesis are likely to be located in the neighborhood (or module) of the five leading edge genes in the functional network. With this in mind, we first constructed a data-driven regulatory gene network based on the transcriptomic data from aged skeletal muscle (**Figure 3A**). WGCNA is a data-driven approach to generate gene-gene co-expression networks from all pairwise correlations, allowing for identification of groups or modules of genes that are highly co-expressed and functionally related^15,43^. In this study, data of transcriptomic responses (log-fold change values) after exercise or immobilization protocols relative to the respective baseline controls were used as input. While the network constructed by WGCNA is commonly referred to as a “*co-expression network*”, given the responsive nature of the network in this study (i.e., the network represents the molecular response to immobilization and exercise), we refer to it as a “*responsive gene regulatory network*” in the following analyses. Using the responsive gene regulatory network, we posited that the mechanistic driver (e.g., transcriptional regulator) of FAPs adipogenesis is in the vicinity of the five leading edge genes we identified (**Figure 3B**).

**Figure 3.**
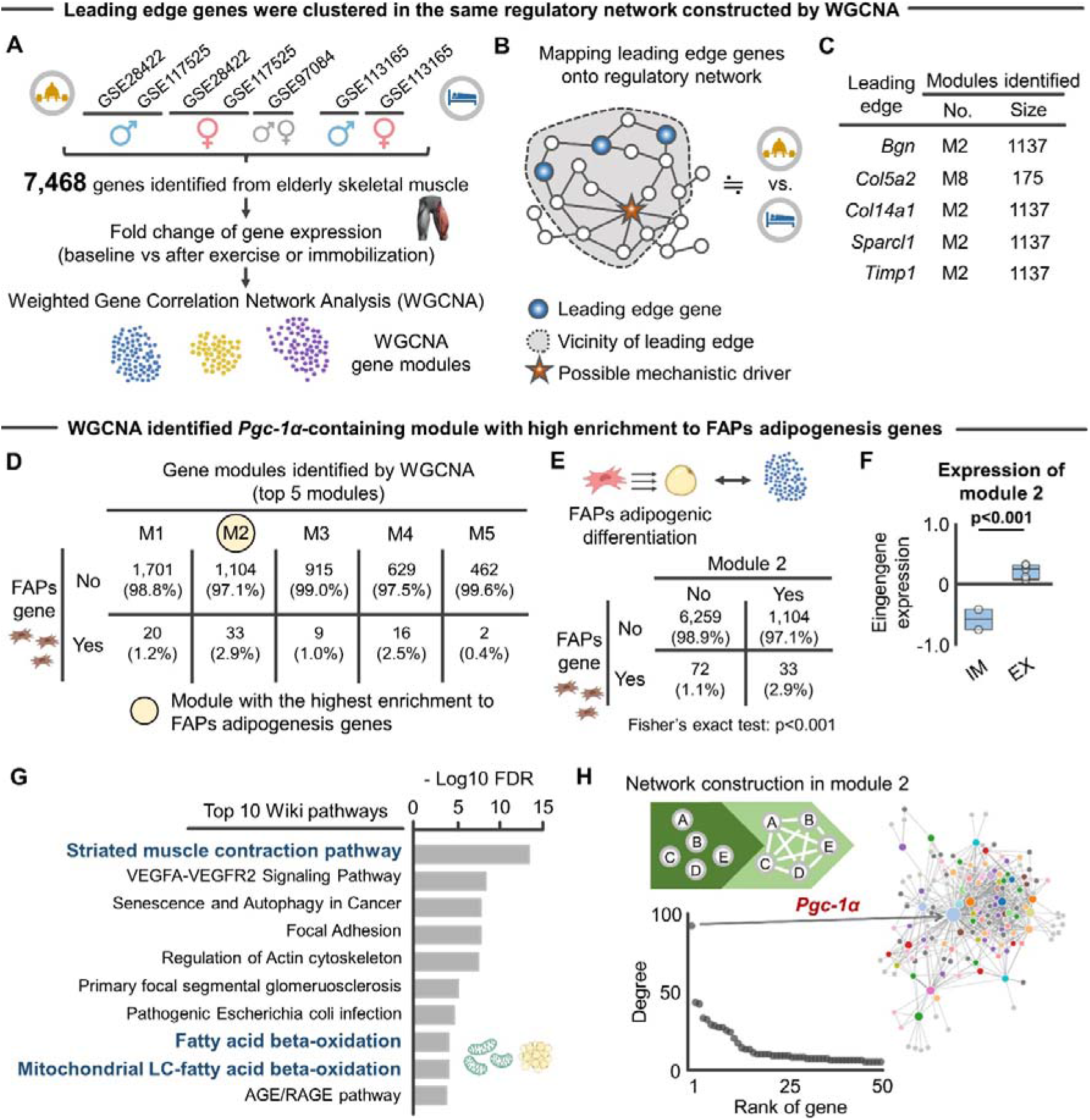
WGCNA revealed co-regulatory relationship between *Pgc-1*α and FAPs adipogenesis. **A**, Schematic showing the analytical flow for WGCNA from the archived transcriptomic data (7,468 genes)^21,25,26,58^. **B**, Schematic showing the concept of identification of mechanistic driver in regulatory network constructed by WGCNA under the assumption of the disease module hypothesis. **C**, Four of the five leading edge genes were included in module 2, the second largest modules constructed. **D-E**, Among the 39 modules, module 2 displayed the highest enrichment to FAPs adipogenesis genes. Values in 2×2 contingency show number of genes (percentage). **F**, Eigengene expression of module 2 was significantly different between immobilization (n = 2) and exercise protocols (n = 5). **G**, Top 10 Wiki pathways significantly enriched to module 2. **H**, Responsive gene regulatory network of module 2. *Pgc-1*α was identified as a hub gene. Statistical analyses were performed using a two-tailed Student *t*-test (**F**) and Fisher’s exact test (**E**). *Abbreviations: EX, exercise; FAPs, fibro-adipogenic progenitors; FDR, false discovery rate; IM, immobilization; WGCNA, weighted gene correlation network analysis*.

Intriguingly, four (*Bgn, Col14a1, Sparcl1*, and *Timp1*) of the five leading-edge genes identified by GSEA were included in module 2, the second largest module (1,137 genes) among the 39 different modules constructed (**Figure 3C**). FAPs adipogenesis genes were similarly highly enriched in module 2 (**Figure 3D-E**). Eigengene expression (a first principal component of a given module) of module 2 was further significantly different across the immobilization and exercise protocols (**Figure 3F**). These findings indicate that genes in module 2 represent FAPs adipogenesis genes that are sensitive to resistance exercise and immobilization, suggesting that module 2 may include mechanistic drivers of FAP adipogenesis. Indeed, Wiki pathway enrichment analysis^44^ identified pathways related to muscle contractile activity and adipogenesis, such as “Striated Muscle Contraction Pathway”, “Fatty acid beta-oxidation”, and “Mitochondrial LC-Fatty Acid Beta-Oxidation” (**Figure 3G**). Most notably, the constructed responsive gene regulatory network identified PPARγ coactivator-1α (*Pgc-1*α) as a hub gene in module 2 (**Figure 3H**). *Pgc-1*α is a transcriptional coactivator known to play a major role in adaptive remodeling of skeletal muscle mitochondria^45,46^. *Pgc-1*α also targets a number of genes involved in lipid droplet assembly and mobilization in response to exercise^47^. Moreover, a recent study has shown that myofiber-specific overexpression of *Pgc-1*α inhibits the accumulation of intramuscular adipocytes in middle-aged mice following glycerol injection^48^. Intriguingly, FAPs in mice with Duchenne muscular dystrophy are characterized by dysfunctional mitochondrial metabolism and increased adipogenic potential, and metabolic reprogramming of mitochondria in FAPs inhibited the adipogenic differentiation^49^. Building upon this previous work, this study identified a large gene module that represent biological function associated with FAPs adipogenesis, in which *Pgc-1*α was implicated as a central regulator.

### GSEA-guided network propagation identified *Pgc-1*α as a possible driver of FAP adipogenesis induced by exercise and immobilization

To further support the finding that *Pgc-1*α is a hub gene in module 2 and, therefore, a possible mechanistic drive of FAPs adipogenesis regulation induced by resistance exercise and immobilization, we herein introduced GSEA-guided network propagation on the responsive gene regulatory network. Network propagation explores the network vicinity of genes seeded to study their biological functions based on the premise that nodes related to similar functions tend to lie close to each other in the networks^50^ (**Figure 4A**). We used a random-walk-restart (RWR) algorithm^51^ to verify which nodes are more frequently visited on a random path in a given network. We posited that RWR starting at the five leading edge genes (i.e., *in silico* over expression) would consistently pseudo activate (i.e., affinity score>0) the *Pgc-1*α gene node (**Figure 4B**). **Figure 4C** shows an example of *in silico* over expression of *Sparcl1* on the network.

**Figure 4.**
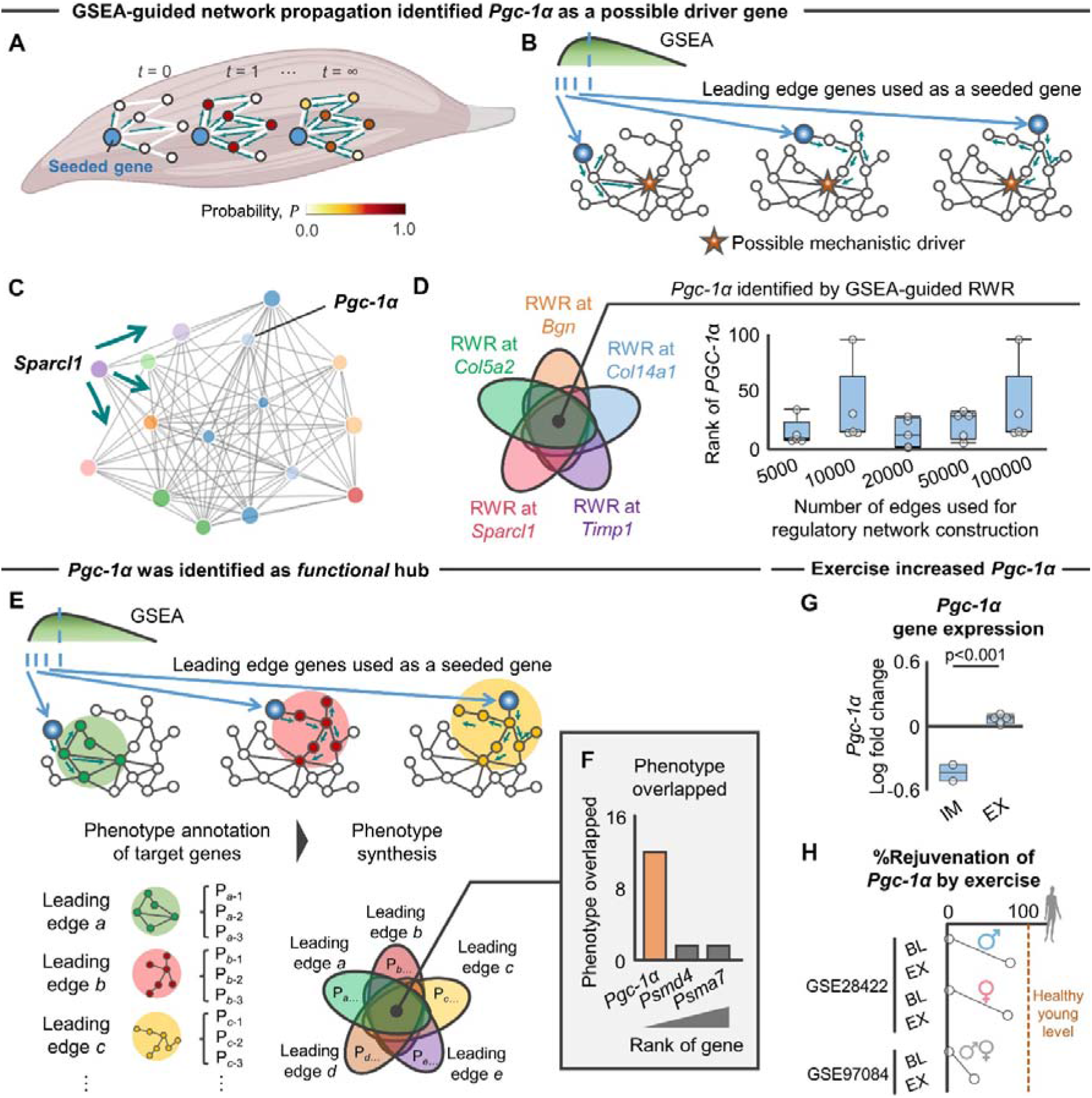
GSEA-guided network propagation identified *Pgc-1*α as a possible driver gene. **A**, Schematic showing the concept of network propagation on the *de novo* responsive gene regulatory network constructed by WGCNA. **B**, Schematic showing the network paradigm of GSEA-guided network propagation starting at the five leading edge genes identified by GSEA. GSEA-guided network propagation implemented multiple RWR with the goal of the identification of possible mechanistic drivers (highlighted as red star) of phenotype associated with the leading edge genes. **C**, Example of RWR started at *Sparcl1*, illustrating 15 vicinity genes of *Sparcl1* that include *Pgc-1*α. **D**, GSEA-guided RWR consistently visited *Pgc-1*α within top 100 rank of the probability score. **E**, Schematic showing GSEA-guided network propagation with subsequent phenotypic (i.e., Wiki pathways) annotation to identify phenotypes associated with the five leading edge genes. **F**, *Pgc-1*α was identified as a functional hub. The graph shows the number of phenotypes overlapping in each gene. **G**, The *Pgc-1*α response was significantly different between immobilization (n = 2) and exercise protocols (n = 5). **H**, %Rejuvenation of *Pgc-1*α by resistance exercise (0%: elderly at baseline, 100%: healthy young adults). Statistical analysis was performed using two-tailed Student *t*-test (**G**). *Abbreviations: EX, exercise; FAPs, fibro-adipogenic progenitors; GSEA, gene set enrichment analysis; IM, immobilization; RWR, random walk with restart; WGCNA, weighted gene correlation network analysis*.

As hypothesized, *in silico* pseudo activation of the five genes consistently pseudo activated *Pgc-1*α gene node (**Figure 4D**). Of note, the responsive gene regulatory network was constructed using the top 5000 co-regulated genes pairs as a default. To address the possibility that different numbers of co-regulated genes pairs for the network construction (i.e., different network topology) may influence the ability of network propagation to identify *Pgc-1*α node, we performed sensitivity analyses in which greater number of co-regulated genes pairs (10,000, 20,000, 50,000, and 100,000 edges) were used for the network construction. Regardless of the varied network topologies, *in silico* pseudo activation of the five genes consistently visited on *Pgc-1*α gene node (**Figure 4D**). This finding further supports the conclusion that the five leading edge genes are co-regulated with the *Pgc-1*α gene following immobilization and resistance exercise.

To further determine whether *Pgc-1*α serves as “functional” hub of the FAP adipogenesis gene module, we repeated the RWR starting at the five leading edge genes and determined phenotypes associated with pseudo activated genes after each network propagation (i.e., significantly enriched Wiki pathways^44^; **Figure 4E**). The *in silico* overexpression of the five leading-edge FAP adipogenesis genes consistently regulated 19 different Wiki pathways, in which *Pgc-1*α was the hub gene contributing to 12 (63.2%) Wiki pathways (**Figure 4F**). Together, GSEA-guided network propagation identified *Pgc-1*α as a functional hub of the responsive regulatory network for FAP adipogenesis.

Since *Pgc-1*α is upregulated by resistance exercise (**Figure 4G**), we also asked whether upregulated *Pgc-1*α gene expression in elderly individuals following resistance exercise approximates the level of healthy young adults. For this purpose, we revisited the MetaMEx database and accessed transcriptomic data of skeletal muscle in healthy young individuals without exercise intervention (2 studies with 3 data sets; mean age: 25.1 years old; body mass index: 24.6 kg/m^2^; n = 29 participants)^18,19^. To account for batch effects induced by inter-trial methodological heterogeneity, we used the transcriptomic data of healthy young participants that was published from the same papers of resistance exercise as the elderly participants (GSE28422 and GSE97084)^18,19^. We then calculated the “*% rejuvenation*” of *Pgc-1*α expression (0%: elderly skeletal muscle without exercise, 100%: healthy young muscle without exercise) induced by resistance exercise. Results showed that *Pgc-1*α expression was recovered to approximately 60% on average relative to healthy young muscle, although we found inter-trial heterogeneity in the *Pgc-1*α response (**Figure 4H**). Of note, consistent with FAPs adipogenesis genes (**Figure 2B**), male and female participants displayed similar transcriptomic response of *Pgc-1*α to resistance exercise (**Figure 2B**). We interpret this data to suggest that resistance exercise has rejuvenating effects on expression of the functional hub gene *Pgc-1*α independent of sex, ultimately driving downstream signals that decrease IMAT accumulation.

### *In silico* and *in situ* activation of *Pgc-1*α targets mitochondrial fatty acid oxidation

The findings shown above taken in the light of previous literature suggest that *Pgc-1*α is an activity-dependent driver gene for FAPs adipogenesis in skeletal muscle. To address the possible downstream signaling of *Pgc-1*α, we again performed RWR that induces *in silico* pseudo activation of the *Pgc-1*α gene and assessed signaling pathways associated with genes in the vicinity of *Pgc-1*α in the reactive gene regulatory network (**Figure 5A**). The RWR started at *Pgc-1*α visited 715 genes, which significantly activated 53 different pathways, including those associated with mitochondria-mediate lipid metabolism such as “Fatty acid beta-oxidation” and “Mitochondrial LC-fatty acid beta-oxidation” (**Figure 5B**).

**Figure 5.**
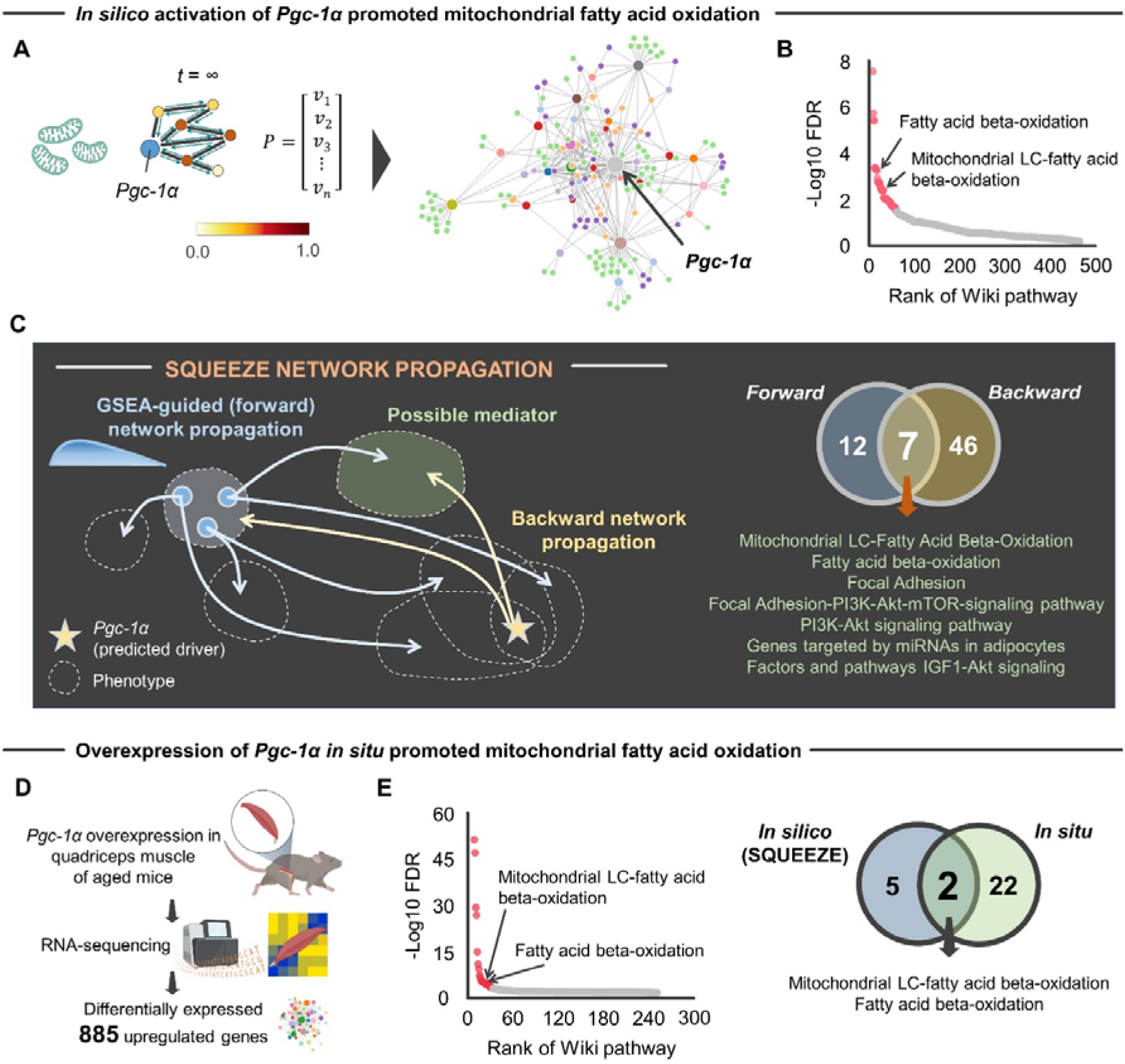
*In silico* and *in situ* activation of *Pgc-1*α targeted mitochondrial fatty acid oxidation. **A**, Schematic showing RWR started at *Pgc-1*α gene node (highlighted by blue color) on the *de novo* gene regulatory network constructed by WGCNA. Gene regulatory network constructed by *Pgc-1*α pseudo activation is provided. **B**, Wiki pathway enrichment analysis of the gene regulatory network constructed by *Pgc-1*α pseudo activation. Pathways significantly (FDR <0.05) modulated were illustrated as red color. **C**, Schematic illustrating the concept of squeeze network propagation, a bidirectional approach integrating GSEA-guided leading edge (forward) and *Pgc-1*α (backward) network propagations. The squeeze network propagation identified seven Wiki pathways which are possible mediators of downstream signal of *Pgc-1*α towards regulation of FAPs adipogenesis. **D**, Schematic showing the experimental flow of RNA-seq after *Pgc-1*α overexpression in the quadriceps muscle of aged mice *in situ*, identifying 885 genes significantly upregulated compared to wild type control aged mice^46^. **E**, Wiki pathway enrichment analysis of the *Pgc-1*α target genes. Pathways significantly (FDR <0.05) modulated were illustrated as red color. Two pathways, “Fatty acid beta-oxidation” and “Mitochondrial LC-fatty acid beta-oxidation”, were consistently modulated by *in silico* and *in situ* activation of *Pgc-1*α *Abbreviation: FAPs, fibro-adipogenic progenitors; FDR, false discovery rate; GSEA, gene set enrichment analysis; RWR, random walk with restart*; *WGCNA, weighted gene correlation network analysis*.

Since the network propagation of *Pgc-1*α activates all pathways in the vicinity of *Pgc-1*α, these pathways do not necessarily mediate the *Pgc-1*α-driven regulation for FAPs adipogenesis. To address this issue, we integrated the two independent network propagations starting at the exercise-responsive five leading edge FAPs adipogenesis genes and *Pgc-1*α (**Figure 5C**). The assumption of this approach is that overlapping pathways of GSEA-guided (forward) network propagation (i.e., starting at FAPs adipogenesis genes) and backward (i.e., starting at *Pgc-1*α) network propagation are possible mediators. This bidirectional network propagation, or “squeeze network propagation”, identified seven pathways, including as “Fatty acid beta-oxidation”, and “Mitochondrial LC-fatty acid beta-oxidation” (**Figure 5C**).

To further validate the *in silico* network inference, we accessed archived RNA-seq data from aged mice with *Pgc-1*α overexpression in quadriceps muscle, which yielded 885 significantly upregulated genes when compared to wild type aged mice^46^ (**Figure 5D**). Consistent with *in silico* prediction (**Figure 5B**), we identified “Fatty acid beta-oxidation”, and “Mitochondrial LC-fatty acid beta-oxidation” as significant pathways associated with the 885 genes upregulated after *Pgc-1*α activation *in situ* (**Figure 5E**). Intriguingly, these two pathways were the only overlapping pathways identified by both *in silico* and *in situ* activation of *Pgc-1*α (**Figure 5E**). These data suggest that these two pathways are mediators of *Pgc-1*α-driven regulation of FAP adipogenesis.

Finally, we asked whether and how resistance exercise and immobilization regulate mitochondrial fatty oxidation. PCA revealed a clear segregation of the transcriptomic response of genes related to mitochondrial fatty acid oxidation when comparing immobilization and resistance exercise protocols (**Figure 6A**). Specifically, we found that these genes were upregulated by resistance exercise but downregulated by immobilization (**Figure 6B**). The responses of these genes to resistance exercise and immobilization protocols were correlated with those of FAPs adipogenesis (**Figure 6C**). Individual gene level analysis revealed that physical activity level was positively associated with expression of key genes involved in mitochondrial fatty acid oxidation (**Figure 6D**). These analyses support the hypothesis that resistance exercise upregulates *Pgc-1*α and promotes mitochondrial fatty acid oxidation to suppress FAPs adipogenesis, but that immobility has the opposite effect.

**Figure 6.**
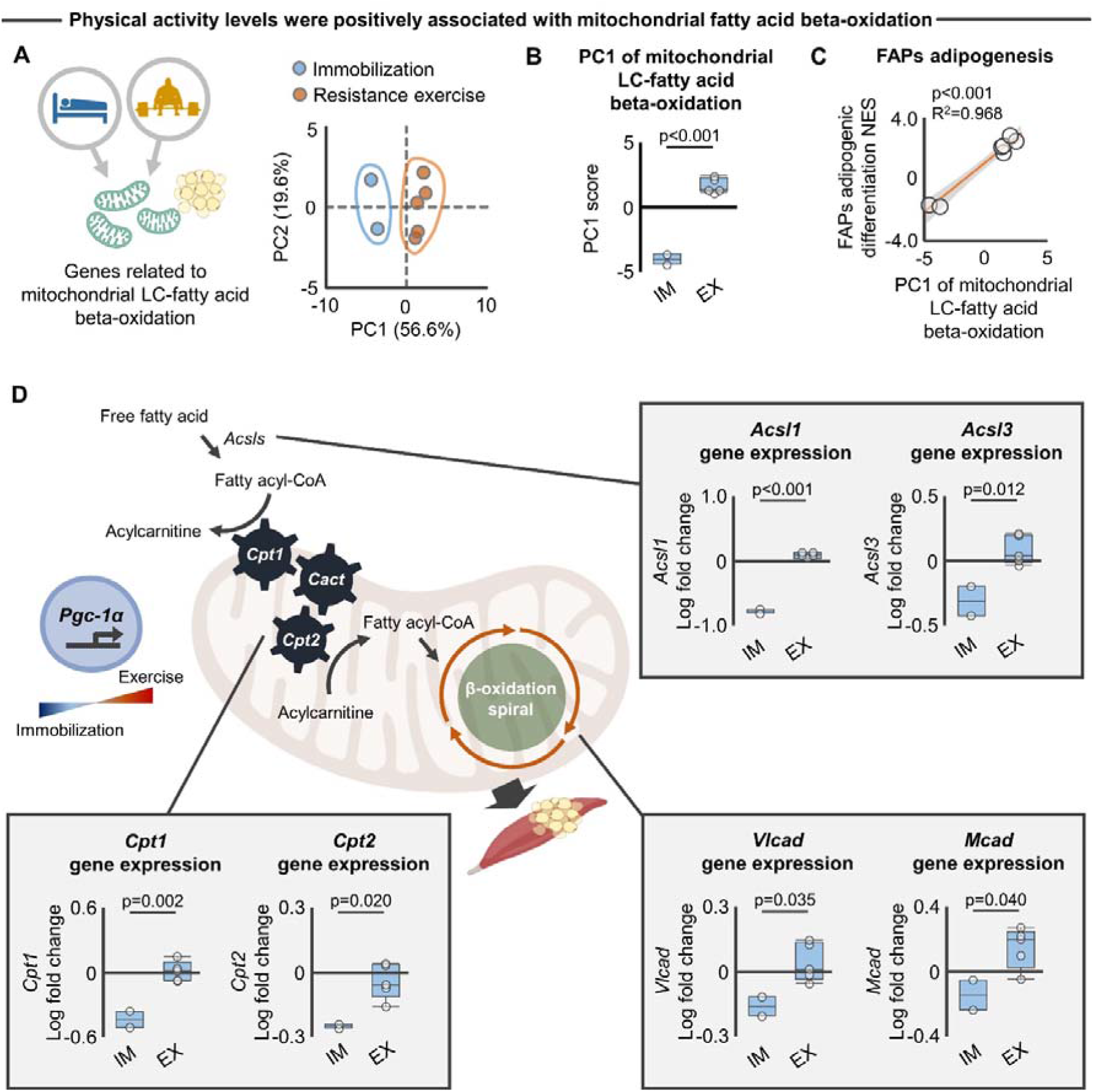
Individual genes response of mitochondrial LC-fatty acid beta-oxidation to exercise and immobilization protocols. **A**, PCA showing the separate clusters in immobilization (blue circle; n = 2) and resistance exercise (orange circle; n = 5) against to overall expression of genes related to mitochondrial LC-fatty acid beta-oxidation. **B**, PC1 extracted from the PCA revealed the significantly different transcriptomic response of the genes related to mitochondrial LC-fatty acid beta-oxidation to immobilization (n = 2) and resistance exercise protocols (n = 5). **C**, Transcriptomic response of the genes related to mitochondrial LC-fatty acid beta-oxidation to immobilization (n = 2) and resistance exercise protocols (n = 5) was significantly associated with regulation of FAPs adipogenesis (i.e., NES score). NES score is calculated by GSEA (see Figure 2B). **D**, Transcriptomic response of key individual genes involved in mitochondrial LC-fatty acid beta-oxidation to immobilization (n = 2) and resistance exercise protocols (n = 5). Statistical analyses were performed using two-tailed Student *t*-test (**B, D**) and linear regression (**C**). *Abbreviations: EX, exercise; FAPs, fibro-adipogenic progenitors; GSEA, gene set enrichment analysis; IM, immobilization; NES, normalized enrichment score; PC, principal component; PCA, principal component analysis*.

## DISCUSSION

Physical exercise has been a widely recognized pillar in the management of skeletal muscle health in older individuals. Accumulating clinical evidence has shown that physical exercise reduces ectopic deposition of fat in skeletal muscle in elderly and patients with chronic diseases^11^. However, the molecular mechanisms underlying the adipose regulating effects of exercise have been poorly understood, thereby representing a critical knowledge gap in the development of effective health-promoting rehabilitative strategies for this population. Here, we took unbiased approach with a series of computational analyses to identify mechanistic drivers of exercise-induced IMAT regulation. A summary of our findings is shown in the **Graphical abstract**. After identifying a set of genes associated with FAP adipogenesis from publicly archived datasets, we discovered that resistance exercise and immobilization in aged skeletal muscle displayed opposing regulatory effects on FAP adipogenesis. Integrated analysis of WGCNA and GSEA-guided network propagation revealed that the exercise- and immobilization-driven regulation on FAP adipogenesis was co-expressed with *Pgc-1*α gene response, where *Pgc-1*α acted as a functional hub for the larger responsive gene regulatory network. *In silico* network inference of squeeze network propagation integrated with *in situ* activation of *Pgc-1*α identified mitochondrial fatty acid oxidation as a mediator of *Pgc-1*α-driven regulation of FAPs adipogenesis. Together, the findings of this current study suggest that physical activity reduces IMAT deposition via upregulation of *Pgc-1*α-mediated mitochondrial fatty acid oxidation and subsequent inhibition of FAP adipogenesis.

The relationship between FAP adipogenesis, a primary inducer of IMAT^5-7^, and chronic physical exercise, a known suppressor of IMAT^11^, has been poorly understood. To address this critical shortcoming, this study identified genes associated with FAP adipogenesis and performed GSEA to demonstrate that chronic resistance exercise and immobilization differentially regulated FAP adipogenesis. This trend was consistent across the different transcriptomic datasets of resistance exercise, indicating the relationship between exercise-responsive genes and FAPs adipogenesis is robust. Since the adipogenic inhibitor, *Sparcl1*, was identified as the primary leading-edge gene upregulated by resistance exercise, we interpreted this finding to suggest that upregulation of *Sparcl1* in skeletal muscle contributes to exercise-induced IMAT inhibition. Interestingly, the regulatory effects of FAP adipogenesis by resistance exercise and immobilization were comparable across the male and female participants. Our results are in line with previous meta-analyses demonstrating that both males and females display a significant reduction in visceral fat after exercise^52^. Nevertheless, there are conflicting results on the effects of sex on the mobilization and oxidation of endogenous triglycerides during exercise^53-55^. Sex-dependent regulation of lipid metabolism by exercise remains an interesting area for future investigation.

To interrogate potential mechanisms underlying the effects of exercise and immobilization on FAP lineage specification, this study introduced the network paradigm, GSEA-guided network propagation, under the assumption of a “disease module hypothesis”^42^, where disease-associated genes or proteins likely share the same topological neighborhood in a network^42^. In this study, we used a GSEA leading edge as a start of network propagation on the WGCNA-based responsive gene regulatory network to explore upstream regulators of exercise-induced FAP adipogenesis. These analyses successfully identified the transcription cofactor gene, *Pgc-1*α, as differentially regulated by immobilization and exercise. *Pgc-1*α has been known as a master regulator of mitochondrial biogenesis in the skeletal muscle response to exercise^56^. Building upon these previous findings, this study suggests a previously unappreciated mechanistic role of *Pgc-1*α in the regulation of FAP adipogenesis and IMAT accumulation in aged skeletal muscle. These findings support the utility of GSEA-guided network propagation as a powerful tool to predict the transcriptional regulator of the skeletal muscle health. This network propagation-based approach is further strengthened by combining another set of network propagation that we first introduced as “squeeze network propagation”. This bidirectional network propagation allows for discovery of novel candidate mediators between the seeded two genes or phenotypes. Through this analysis, we successfully prioritized possible pathways and identified mitochondrial fatty oxidation as a primary driver for FAP regulation by *Pgc-1*α.

Although this study provides novel insight to the pathogenesis of age-related IMAT accumulation, it has limitations. First, findings of this study are based on a small number of datasets, particularly in the context of immobilization. This may contribute to bias depending on the participants’ characteristics and the methodology used in the original studies. Another limitation is that the possible working mechanism proposed in this study is based on transcript levels only. An important next step will be to interrogate the identified mechanism at the protein and tissue levels.

While this study focused on skeletal muscle, a major conceptual innovation of the proposed method is the demonstration of the mechanistic link between exercise-driven health promoting effects and target disease (i.e., IMAT in elderly). It is widely recognized that physical exercise elicits systemic benefits beyond skeletal muscle in both young and older populations^57^. Yet, the mechanisms of the multi-systemic benefits induced by exercise remain largely unknown. We anticipate that the network paradigm proposed in this study may have broader implications in the field of exercise physiology and aging research.

## Acknowledgments

This study was supported in part by Tokai Pathways to Global Excellence Project (T-GEx) (https://www.t-gex.nagoya-u.ac.jp/en/). The funders had no role in study design, data collection and analysis, decision to publish, or preparation of the manuscript.

## Author contributions

All authors made substantial contributions in the following areas: (1) conception and design of the study, acquisition of data, analysis and interpretation of data, drafting of the article; (2) final approval of the article version to be submitted; and (3) agreement to be personally accountable for the author’s own contributions and to ensure that questions related to the accuracy are appropriately investigated, resolved, and the resolution documented in the literature.

The specific contributions of the authors are as follows:

H.I., F.A., Y.M. provided the concept, idea and experimental design for the studies. H.I., F.A., Y.M. wrote the manuscript. H.I., F.A., Y.M. provided data collection, analyses, interpretation and review of the manuscript. H.I. obtained funding for the studies.

## Inclusion and diversity

We support inclusive, diverse, and equitable conduct of research.

## STAR METHODS

### KEY RESOURCE TABLE

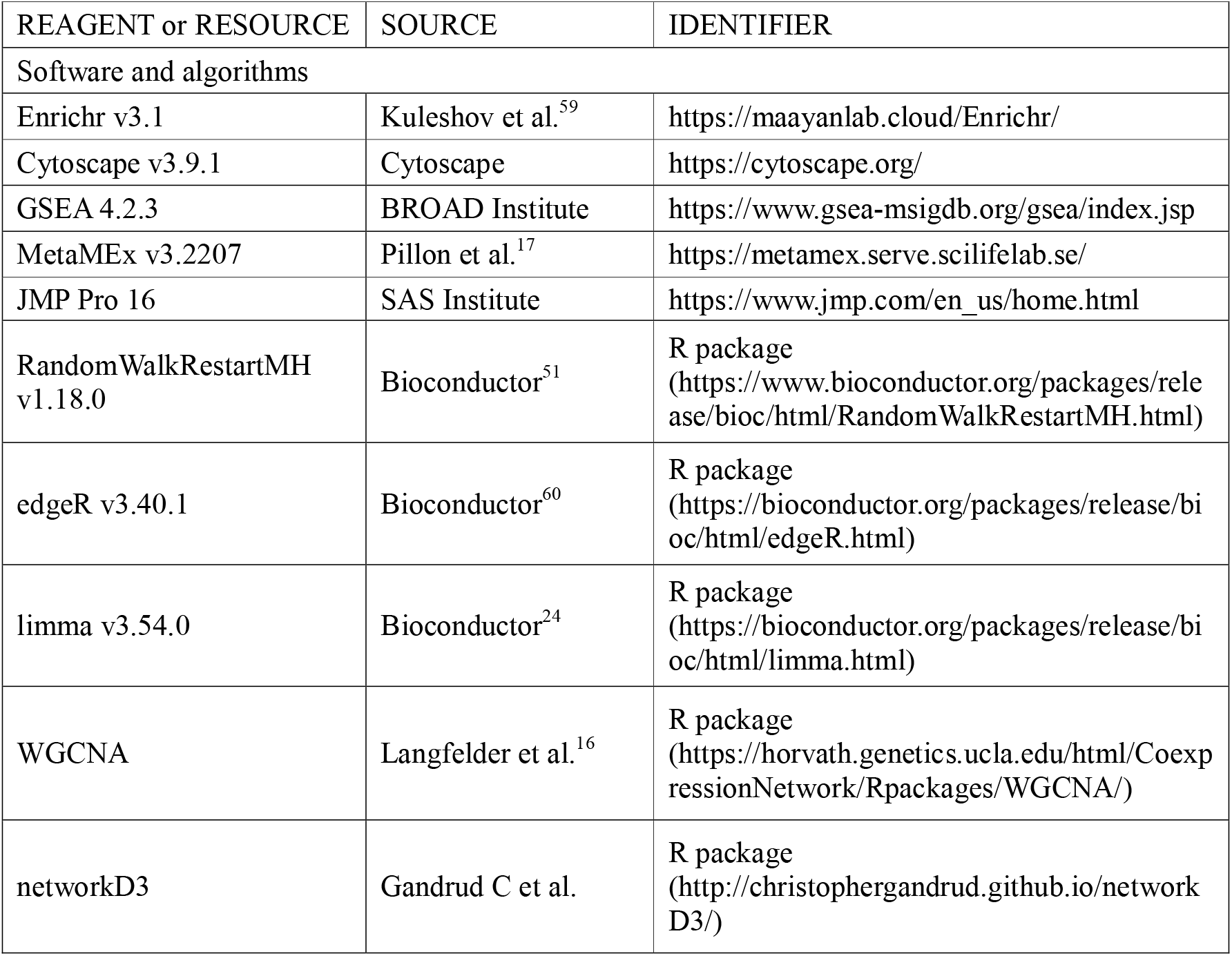

| REAGENT or RESOURCE | SOURCE | IDENTIFIER |
| --- | --- | --- |
| Software and algorithms |  |  |
| Enrichr v3.1 | Kuleshov et al.<sup>59</sup> | https://maayanlab.cloud/Enrichr/ |
| Cytoscape v3.9.1 | Cytoscape | https://cytoscape.org/ |
| GSEA 4.2.3 | BROAD Institute | https://www.gsea-msigdb.org/gsea/index.jsp |
| MetaMEx v3.2207 | Pillon et al.<sup>17</sup> | https://metamex.serve.scilifelab.se/ |
| JMP Pro 16 | SAS Institute | https://www.jmp.com/en_us/home.html |
| RandomWalkRestartMH v1.18.0 | Bioconductor<sup>51</sup> | R package<br>(https://www.bioconductor.org/packages/release/bioc/html/RandomWalkRestartMH.html) |
| edgeR v3.40.1 | Bioconductor<sup>60</sup> | R package<br>(https://bioconductor.org/packages/release/bioc/html/edgeR.html) |
| limma v3.54.0 | Bioconductor<sup>24</sup> | R package<br>(https://bioconductor.org/packages/release/bioc/html/limma.html) |
| WGCNA | Langfelder et al.<sup>16</sup> | R package<br>(https://horvath.genetics.ucla.edu/html/CoexpressionNetwork/Rpackages/WGCNA/) |
| networkD3 | Gandrud C et al. | R package<br>(http://christophergandrud.github.io/networkD3/) |

## RESOURCE AVAILABILITY

### Lead contact

Further information and requests for resources should be directed to and will be fulfilled by the lead contact, Hirotaka Iijima.

### Materials availability

This study did not generate new unique reagents.

### Data and code availability

The raw data that support the experimental findings are included as Source Data. Any additional information required to reanalyze the data reported in this work is available from the lead author upon request.

## EXPERIMENTAL MODEL AND SUBJECT DETAILS

### Human subjects

This study used archived data of human experiments.

### Animals

This study used archived data of animal experiments.

## METHODOLOGICAL DETAILS

### Search for transcriptomic data of elderly skeletal muscle response to exercise and immobilization

We accessed publicly available transcriptomic data of skeletal muscle generated by MetaMEx project^17^. MetaMEx summarizes skeletal muscle transcriptomic response to immobilization and exercise (aerobic, resistance, and high intensity training) across the lifespan of human adults (young, middle, and elderly)^17^. We first accessed all the raw data through GitHub (https://github.com/NicoPillon/MetaMEx). According to the pre-determined eligibility criteria, we included studies assessing the transcriptomic response to immobilization or chronic (>3months) resistance exercise protocols in healthy elderly participants. Elderly was defined as age over 65 years in each study. No restrictions were set according to target muscle of biopsy, quantification method for skeletal muscle transcripts, participants’ sex, body mass, race, country, and publication year. For the data sets of resistance exercise, this study limited participants with physical activity level of sedentary because of possible confounding effects on the transcriptomic response to exercise protocol^61^. The MetaMEx database search was performed in November, 2022 by single reviewer (HI). During these processes, the reviewer prepared and used simple, pre-designed Google spreadsheets to assess eligibility by extracting study features.

### Identification of genes associated with insulin sensitivity in elderly skeletal muscle

Genes associated with insulin sensitivity were defined according to analyzed data in previous literature^27^. The original study published by Lutter et al^27^. obtained transcriptomic profile as well as clinical data of insulin sensitivity (e.g., glucose infusion rate) from skeletal muscle and IMAT in 16 elderly participants with obesity and type II diabetes mellitus from a cross-sectional study previously published by Sachs et al^2^. After multivariate regression analysis in which gene expression were regressed to insulin sensitivity, Lutter et al identified 59 genes positively associated with good insulin sensitivity^27^. We have accessed the list of genes and used for further analyses.

### Mouse model of skeletal muscle-specific overexpression of Pgc-1α gene

We accessed the archived RNA-seq data collected from quadriceps muscle in older-old (29-34 months old) male and female C57BL/6J mice^46^. The original study published by Garcia et al identified 1,485 differentially regulated genes (false discovery rate <0.05, fold change >2), of which 885 and 600 were up- and downregulated, respectively^46^. Since the transcriptomic response was similar across males and females, the RNA-seq data from males and females were mixed and used for further analyses.

## QUANTIFICATION AND STATISTICAL ANALYSIS

### RNA-seq analysis to define genes associated with FAPs adipogenesis

We accessed the archived RNA-seq data (GSE171502) collected from isolated FAPs with and without miR-206 deficiency (as induced by siRNA to miR-206)^33^. The miR-206 deficiency recapitulated increased adipogenic differentiation of PDGFRα-positive FAPs *in vitro*^33^. Raw count data was normalized by count per million (CPM), filterByExpr function, and Trimmed Mean Mvalue (TMM) using R/Bioconductor package edgeR with default parameters^60^. Differential gene expression analysis was performed for genes with a normalized CPM value using R/Bioconductor package limma^24^. The Benjamini-Hochberg FDR control for multiple hypothesis testing was used to produce q-values.

### Filtering low expression genes in MetaMEx transcriptomic data

We removed genes that have very low counts in RNA-seq or signal intensity in microarray data prior to downstream analysis from biological and statistical grounds^22^. From biological point of view, a gene must be expressed at some minimal level before it is likely to be translated into a protein or to be considered biologically important. From a statistical point of view, genes with consistently low counts or low signal intensity are very unlikely be assessed as differentially express because low counts or low signal intensity do not provide enough statistical evidence for a reliable judgement to be made. Such genes can therefore be removed from the analysis without any loss of information. We used filterByExpr function for count data in RNA-seq data^22^. For signal intensity in microarray data, we performed soft intensity based filter^62^ as recommended by R/Bioconductor package limma user guide^24^.

### Functional characterization of transcriptome using pathway enrichment analysis

To determine the biological function of genes of interests, Wiki pathway enrichment analysis was performed by Enrichr software^59^. WikiPathways (wikipathways.org) captures the collective knowledge represented in biological pathways^44^. WikiPathways provides easy-to-use drawing and annotation tools to capture identities, relationships, comments and literature references for each pathway element and interaction.

### GSEA

GSEA was performed as described previously^35^ using the GSEA web tool provided by Broad Institute Website (https://www.gsea-msigdb.org/gsea/index.jsp). This study implemented ssGSEA^63^, an extension of GSEA that allows one to define an enrichment score that represents the degree of absolute enrichment of a gene set in each sample within a given data set. The transcriptomic responses (i.e., log fold change) of elderly skeletal muscle to resistance exercise and immobilization relative to baseline control were rank-normalized and used for input. The genes associated with insulin sensitivity or FAPs adipogenesis we originally defined were used as a gene set. The minimum and maximum criteria for selection of gene sets from the collection were 10 and 500 genes, respectively.

Leading edge analysis was performed after each GSEA to determine the core genes define the subset of genes with positive contribution to the enrichment score before it reaches its peak; i.e., those that are most correlated with the phenotype of interest. These leading edge genes were used as seeded genes for network propagation with RWR.

### WGCNA and hub gene analysis

This study used the WGCNA package to build a weighted gene co-expression network using the archived RNA-seq or microarray data in MetaMEx database, which finally yielded 7,468 genes identified across the different data sets after filtering low expression genes, as described above. The key parameter, β, for weighted network construction was optimized to maintain both the scale-free topology and sufficient node connectivity as recommended in the manual. A topological overlap matrix (TOM) was then formulated based on the adjacency values to calculate the corresponding dissimilarity (1-TOM) values. Module identification was accomplished with the dynamic tree cut method by hierarchically clustering genes using 1-TOM as the distance measure with a minimum size cutoff of 30 and a deep split value of 2 for the resulting dendrogram. A module preservation function was used to verify the stability of the identified modules by calculating module preservation and quality statistics in the WGCNA package^16^.

From the modules constructed by WGCNA, we identified hub gene via Cytoscape software (version 3.9.1) using “analyze network” option which determines the topological parameters (e.g., EdgeCount and NeighborhoodConnectivity) of given networks. In the provided parameters, “EdgeCount” describes the link between one node with their adjacent nodes and the node with the highest number of “EdgeCount” was defined as the hub gene in the network.

### Network propagation using RWR

RWR was performed by R/Bioconductor package RandomWalkRestartMH^51^. RWR simulated a walker starting from one node or a set of nodes (seed nodes) in one network, and such walker randomly moved in the network to deliver probabilities on the seed nodes to other nodes. After iteratively reaching stability, the affinity score of all nodes in the given network to seeded node were obtained.

### Quantification of %rejuvenation of Pgc-1α induced by resistance exercise

We accessed two data sets (GSE28422 and GSE97084)^18,19^ of skeletal muscle in (1) healthy young without exercise intervention (at baseline), (2) elderly without exercise intervention (at baseline), and (3) elderly with resistance exercise. From *Pgc-1*α gene expression data across the three conditions, we calculated %rejuvenation of *Pgc-1*α expression (0%: elderly skeletal muscle without exercise, 100%: healthy young muscle without exercise).

### Unsupervised machine learning

PCA was performed for data reduction to identify the principal components that represent differences in the given data sets using JMP Pro 16 software (SAS Institute, Cary, NC). PCA produces linear combinations of the original variables to generate the principal components^64^, and visualization is generated by projecting the original data to the first two principal components.

### Statistical analysis

All statistical analyses were performed using JMP Pro 16 software (SAS Institute, Cary, NC). Except where indicated, data are displayed as means, with uncertainty expressed as 95% confidence intervals (mean ± 95% CI). Two-tailed Student t-test, linear regression analysis, or Fisher’s exact test were performed. We checked the features of the regression model by comparing the residuals vs. fitted values (i.e., the residuals had to be normally distributed around zero) and independence between observations. No correction was applied for multiple comparison because outcomes were determined a priori and were highly correlated. No statistical analyses included confounders (e.g., mean body mass in each study or data set) due to the small sample size. We conducted a complete-case analysis in the case of missing data (e.g., genes under detection by RNA-seq). In all experiments, p-values <0.05 were considered statistically significant. Throughout this text, “n” represents the number of independent observations of study or data set. Specific data representation details and statistical procedures are also indicated in the figure legends.

